# Multiply Perturbed Response to Disclose Allosteric Control of Conformational Change: Application to Fluorescent Biosensor Design

**DOI:** 10.1101/2025.02.01.636023

**Authors:** Melike Berksoz, Ali Rana Atilgan, Burak Kocuk, Canan Atilgan

## Abstract

Proteins exhibit remarkable conformational flexibility, enabling precise functional regulation through allostery. A key application of allostery is in the design of protein-based sensors, which detect environmental changes—such as ligand binding or post-translational modifications—and convert these cues into measurable signals (e.g., fluorescence). Here, we investigate a series of ligand-binding proteins that serve as sensing domains in direct-response fluorescent biosensors, wherein ligand binding enhances fluorescence output. We employ a multiple force application approach which we term Multiply Perturbed Response (MPR) to identify “hot spot” residues that drive the conformational transition from an *apo* (inactive/OFF) to a *holo* (active/ON) state. We first present two efficient computational approaches to determine residues and forces that maximize the overlap of the observed conformational change. We then determine the overlap maximizer residues for up to five force insertion locations, and we compare them with actual insertion sites used in existing biosensors. This work enhances the utility of linear response theory-based methods in uncovering multiple functionally significant regions that trigger a known conformational change. The approach might prove useful not only in the design of biosensors, or in locating allosteric sites, but may also find applications in offering physics-based collective variables in mapping conformational transition pathways of proteins.

## Introduction

Proteins are inherently dynamic, adopting multiple conformational states that are crucial for their function.^1–4^ This interdependence of conformational states enables the precise control of protein function, allowing cells to rapidly respond to changes in their environment.^5–8^ A fundamental mechanism of adjusting the conformational landscape of proteins is through allostery. In a typical scenario, ligand binding affects the dynamics of a distant site, thereby leading to shifts in the conformational landscape and driving the equilibrium from an inactive state towards an active state. A classical application of allostery is found in protein-based sensors.^9,10^ These reagents sense shifts in their environment—such as changes in pH, ligand binding events, or post-translational modifications—and respond through conformational reorganization. This rearrangement then conveys a signal to a reporter domain, resulting in outputs that can include enzymatic activation, gene regulation, or fluorescence. Naturally occurring ligand binding proteins have been largely employed as sensing domains in protein biosensors.^11–16^ For an efficient sensor, two conditions must be met; (i) there should be a large enough conformational change upon ligand binding in the sensing domain, (ii) there should be a strong structural coupling between the sensing and the effector domains so that a response to ligand binding can be measured. For example, shifts in hydrogen bonding patterns as a result of *apo*-to-*holo* conformational transition was proposed as a means of allosteric communication between the sensing and reporter domains.^17^ In sensor construction, reporter domains are commonly inserted to an allosteric position distant from the ligand binding site. Insertion sites are often determined by comparing *apo* and *holo* structures of the sensing domains, pinpointing flexible regions with significant backbone dihedral torsion or root mean squared fluctuation (RMSF) changes in MD simulations. In case of Förster Resonance Energy Transfer (FRET) sensors, a more straightforward approach is taken, where two fluorescent proteins (FPs) are attached to the N and C termini of the sensing domain. Change in the end-to-end distance as a result of ligand binding causes the two FPs coming into a suitable distance (<10 nm) for FRET to occur. While there is a lot of information accumulated on the design of genetically encoded fluorescent biosensors (GEFBs) and a plethora of working examples, only a handful of the designs have been obtained from a knowledge of allosteric control in the sensing domains.^11^ In fact, the field is dominated by a trial-and-error approach to the problem by employing high throughput screening of circularly permuted FPs (cpFPs) attached to heuristically selected locations by a variety of viable linkers. ^9,18,19^

Perturbation Response Scanning (PRS) is a popular tool used for the prediction of allosteric sites. In this method, proteins are coarse-grained such that each residue is represented by its C_α_ atom, and each C_α_ atom is systematically perturbed in many directions and the updated coordinates for each force perturbation direction are recorded.^20^ By measuring the overlap between the new coordinates and the experimentally known structures, PRS helps pinpoint protein regions that mechanically modulate binding-site motions, making it particularly useful for identifying functionally significant residues involved in a specific conformational transition. Previously, single residue perturbations were performed on a series of *apo*/*holo* protein pairs with diverse modes of motions, such as hinge bending, or shear.^21^ That study showed that residues whose perturbation induces global displacements aligning with experimental data are indeed functional hot spots, confirmed by both experimental and computational evidence. For example, PRS analysis of the Hsp70 chaperone identified two types of residues: ‘sensors’, which are highly sensitive to allosteric signals, and ‘effectors’, which play a key role in transmitting these signals.^22^ Combined with coevolution analysis, PRS has been used to predict mutational sites of post-translational modifications in heat shock proteins ^22–26^ and mutational hotspots in tumor suppressor proteins.^23^ PRS has been useful in pinpointing allosteric network effects in PDZ domains,^27^ SARS-CoV-2 spike proteins,^28^ β-lactamases^29^ and transferrin,^30^ to name a few. PRS is implemented in the ProDy^31^ and MD-TASK^32^ Python packages, which are widely used in the community.

While exhaustive scanning of a protein for single residue perturbations is a fast process, extending it to multiple site perturbations requires exponentially increasing computational power. An alternative approach is to define this as an optimization problem whereby residue combinations and perturbation directions maximizing the overlap between the positional rearrangements between the *apo*/*holo* forms are sought. In this work, we describe two computational approaches leading to this result, one efficient when a small number of forces are inserted, and the other getting efficient when solutions which require perturbations on larger than four positions are required. We collectively call these approaches Multiply Perturbed Response (MPR). Although the methodology is useful in a range of problems associated with allostery, in this work we use several proteins utilized in the sensor domains of GEFBs to demonstrate outputs and the utility of MPR. We validate our approach by comparing the overlap maximizer residues with sensor insertion sites used in reported fluorescent biosensors.

## Methods

### Basis of the PRS approach

The PRS methodology was first introduced and detailed in ref ^20^. In summary, PRS takes two distinct structural forms of a protein of *N* residues as inputs, referred to as the initial (I) and target (T) structures. The goal is to identify a perturbed residue and a perturbation direction in structure I that produces displacements closely resembling the conformational changes observed between I and T. Using the principles of linear response theory,^33^ PRS establishes a relationship between the applied external force (Δ**F**) and the resulting displacements (Δ**R**) through a 3*N*×3*N* covariance matrix, **C**. The latter is derived from molecular dynamics (MD) simulations or is approximated by the inverse Hessian (**C** ≈ **H**^-1^) constructed from an elastic network model.^34^ The coordinate shifts are computed via,

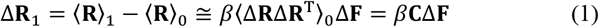

where **R**_0_ and **R**_1_ are the 3*N*×1 vectors with the initial (unperturbed state) and the predicted (perturbed state) C_α_ coordinates of the protein, respectively, and *β* = 1/*k*_*B*_*T*. Δ**F** vector contains the coordinates of the external forces inserted on the residues.

The coarse-grained representation of a state is constructed by taking the C_α_ atom of each residue as a node. Then, by inserting many random, fictitious forces (Δ**F**) in different directions on each node (residue C_α_) to perturb the structure, the predicted Δ**R**_1_ values for every Δ**F** are recorded. These resulting displacements are compared with the conformational change determined from the difference between best-fitted initial and target structures, **R** = **R**_T_ **- R**_I_. As a measure of goodness of prediction, the overlap *O*^*i*^ between the predicted and measured directions is evaluated for the *i*^th^ perturbation:

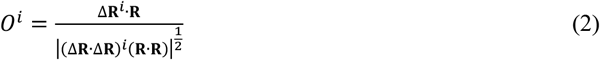

The higher *O*^*i*^ values indicate more similarity to the displacement vector **R** and implies a good choice for perturbing vectors (force vectors). In PRS we seek *O*_max_ which is the maximum amongst all perturbations, after the insertion of *P* perturbations on *N* residues.

### Preparation of PDB files

We list the systems studied in this work in **Table 1**. Each system required various preprocessing to generate the coordinates that are utilized to determine the allosteric regions. These systems were selected from a series of ligand binding proteins which are used as the sensory domains of GEFBs, and they display varying degrees of conformational change as indicated by the RMSD value listed in **Table 1**. Among these, some had protein data bank (PDB) coordinates for either or both *apo* and *holo* forms. For others, we generated coordinates either via the AlphaFold3 (AF3) server^35^ or Colabfold (AF2) with custom settings.^36^

**Table 1.**
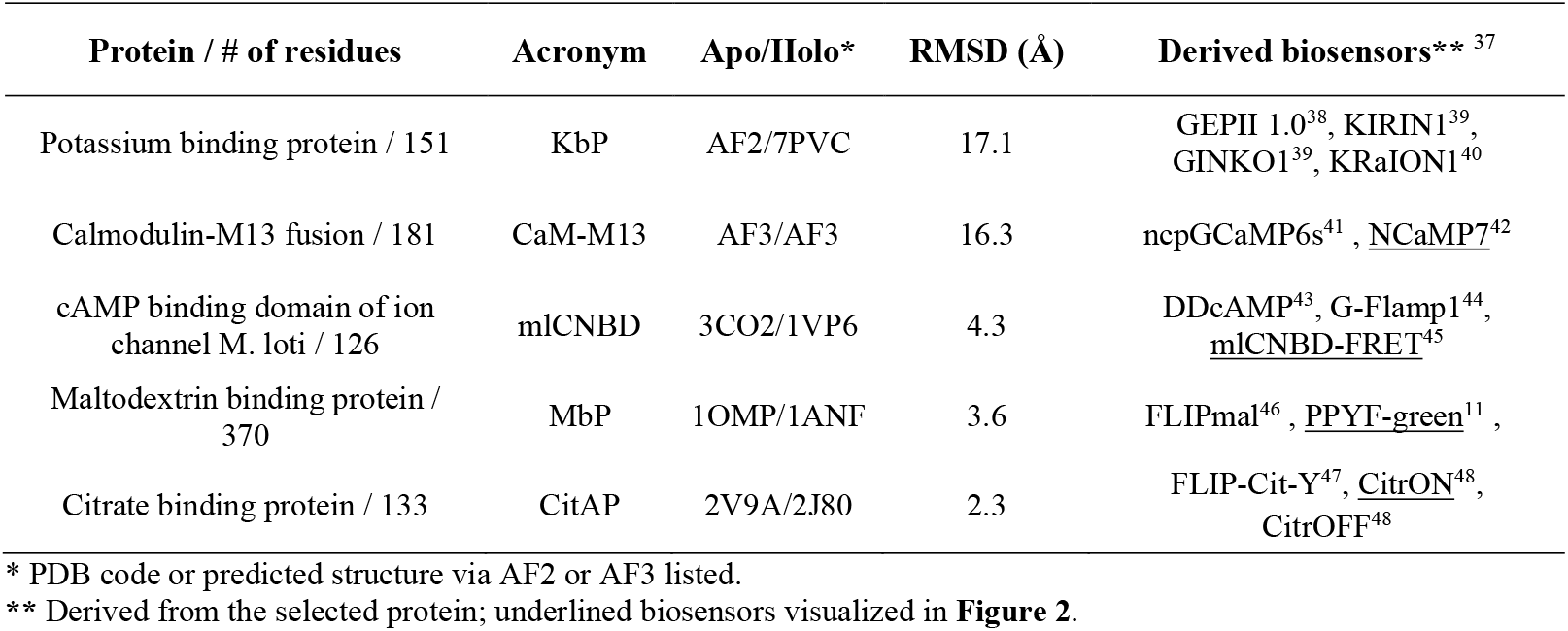
*apo*/*holo* pairs analyzed and the biosensors where the protein is used as the sensing domain.

In case of KbP, AF3 server generated all *holo*-like models, while Colabfold utilizing AF2 with reduced MSA (max_MSA 64:128) could generate one *apo*-like model (**Figure S1A**). The obtained model corresponds well to the elongated structure observed in K^+^-free solution SAXS measurements for this protein.^49^ However, the N terminal β strand and the binding loop, covering residues 1-18, is modelled incorrectly as a long helix. Therefore, we tested a shallower MSA (max_MSA 32:64), which produced all *apo*-like models, with correct modelling of the binding site in the Rank1 model. In the case of CaM-M13 fusion system, the sequence extracted from the sensor NCaMP7 ^42^ was used as input in the AF3 server with the default settings.^35^ The first two ranked models represent the calcium bound form of CaM-M13 domain in NCaMP7, whereas Rank3/4/5 resembles standalone CaM (without the peptide) where the two lobes of CaM are separated with an elongated central α helix (Cα RMSD of 5.4 Å to PDB: 3CLN). In these models, M13 peptide is also segregated from the two lobes **(Figure S1B)**. Although there is no known structure of *apo* CaM-M13, we took two-calcium bound structure of GCaMP2, an earlier version of a CaM-based biosensor, as a reference for an *apo*-like state of CaM-M13.^50^ This structure (PDB: 3O77) shows that the two lobes are separated, with the M13 peptide interacting with the lobe carrying the two calcium ions. Additionally, the central α-helix is bent rather than elongated, as seen in our models. Notably, in this sensor, the M13 peptide is not covalently attached to CaM. Despite these differences, our models bear similarity to this structure in terms of the relative organization of the N- and C-terminal domains, with a Cα RMSD of 10.7 for the CaM domains compared to PDB: 3O77. Both *apo* KbP and *apo* CaM-M13 were predicted with relatively low accuracy as indicated by the color code in Figure S2. However, the closeness to the experimentally known *apo* structures assured us to go forward with these predicted models, following our success using this approach in our previous work.^51^ Structure of mlCNBD-FRET sensor which is comprised of cAMP binding protein mlCNBD and FPs Citrine and Cerulean was predicted with AF3 using the sequences in the fluorescent protein database.^52^ Colabfold with reduced MSA (max_MSA 32:64) was used to complete the missing loop (residues 68-89) in the *apo* structure of citrate binding protein (**Figure S1C**).

When using experimental PDB structures, all sequences were renumbered to start with 1, in case a different numbering was present. All *holo* structures were aligned onto the *apo* structures (unless otherwise stated) using the Biopython^53^ superimposer function and the displacement vectors between the two states were calculated. We note that *apo* and *holo* models should contain the same number of Cα atoms for the calculation of displacement. The Hessian matrix was constructed with a cut-off distance of 12 Å for all systems based on a previous study that scrutinized the effect of the cutoff distance on the converging properties of the essential contributors to **H**.^54^ Morph movies showing the *apo*-to-*holo* conformational transitions were created with the ‘Morph’ command in ChimeraX using the default settings.^55^

### Multiple force application

While PRS reproduces the single residues that maximize *O*^*i*^ in a reasonable amount of computational time, the required time that satisfies the same criterion increases exponentially for multiple forces. We therefore propose two approaches to overcome this shortcoming: (A) via enumeration, and (B) via optimization.

#### A. Enumeration approach

As defined earlier, the difference between the *holo* and *apo* coordinates of a protein with respect to a reference frame is denoted as **R**. Here **R** ∈ *R*^3*N*^ is a nonzero real vector. Let us call **B**_*i*_ = (3*i* − 2,3*i* − 1,3*i*) for *i* = 1, …, *N* as the *i* -th *block*. The linear response theory dictates that

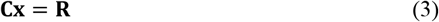

where **x** = *β*Δ**F** in equation 1. Equation 3 has a unique solution **x**^∗^ = **C**^**−1**^**R**. We are interested in the *special* sparse solutions to this system in which a small number of blocks have nonzero entries that have a *large* overlap with **x**^∗^. More formally, we are interested in solving the following optimization problem

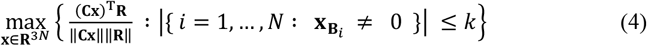

where 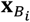 is a three-dimensional vector with entries (*x*_3*i*−2_3 *x*_3*i***−1**_3 *x*_3*i*_) and *‖ ‖* is the Euclidean norm. *k* is the number of atoms on which forces will be inserted in the solution.

##### Solution for a given index set

We first consider the following version of problem (4) for *given* indices *B* ⊆ {1, …, 3*N*} without the consideration of blocks:

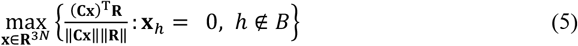

By plugging in **x** = ∑_*h*∈*B*_ α_*h*_**e**_*h*_, where **e**_*h*_ is the *h*-th standard unit vector in *R*^𝟛*N*^, we reformulate problem (5) as

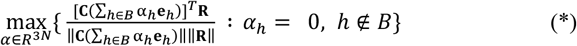

or equivalently as,

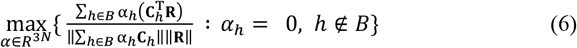

where **C**_*h*_ is the *h*-th column of **C**.

We can find the optimality conditions for problem (6) as

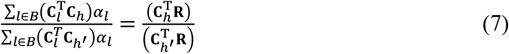

for *h, h*^*′*^ ∈ *B*. Since α values can always be scaled by nonzero scalars, the unique solution of the following linear system satisfies the optimality conditions (7):

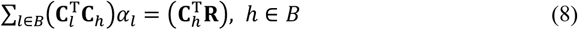

Hence, an optimal solution to problem (5) can be obtained by solving system (8). Coincidentally, this system is the optimality conditions for the following least squares problem in α:

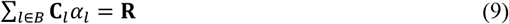

##### Solution for k-block

We will utilize the solution of problem (5) to attack problem (4) with *k* blocks. In fact, we can reformulate problem (4) as

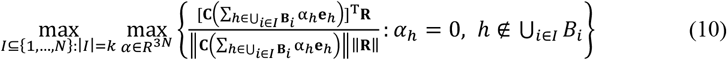

Notice that the inner maximization is an instance of problem (*) with **B** = ⋃_*i*∈*I*_ **B**_*i*_. Therefore, the optimality conditions for the inner maximization can be written as the linear system (8) for **B** = ⋃_*i*∈*I*_ **B**_*i*_. Let us denote the unique solution to this system as α^*I*^. Then, we are ready to solve the outer maximization in problem (10) as,

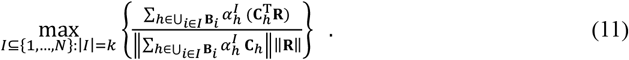

Notice that solving problem (4) via enumeration entails solving 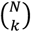 many least squares problems, each with a system of size 3*N* × 3*k*.

#### B. Optimization approach

The optimization approach relies on the fact that the optimality conditions of problem (4) for a given collection of blocks ∪ **B**_*i*_ is exactly the optimality conditions of a least squares problem. Then, we can formulate an optimization problem that seeks the best selection of blocks. The proposed mixed-integer quadratic programming (MIQP) formulation is as follows:

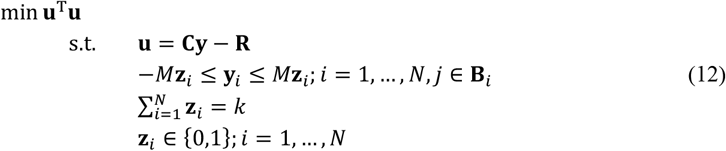

Here, **y** ∈ *R*^𝟛*N*^ is a sparse approximation of **x**^∗^, **u** is the error in the sparse least squares problem and **z**_*i*_ is a binary variable indicating whether block *i* is chosen or not.

In this model, the selection of *M* is very important. This parameter should be chosen large enough so that constraints in equation 12 do not exclude any optimal solutions. However, the value of *M* should not be too large to prevent weak relaxations. Unfortunately, deciding the value of *M* a priori is not very easy. Therefore, we propose to use a simple practical approach: If *k* is small (such as *k* ≤ 3), then it is likely that the enumeration approach would be faster; and if *k* is not very small (such as *k* > 3), it is possible that the optimization approach would be faster. In this case, we propose to first solve *k* = 1, 2, 3 instances with the enumeration approach, record the absolute value of the largest coefficient observed as *M* and use it in the optimization approach for larger values of *k*.

## Results and Discussion

Single residue perturbations have been used to identify key residues involved in the conformational transitions of a variety of proteins. Although PRS is based on a network of Cα atoms, the character of these residues is important as indicated by their high level of conservation or mutations that compromise the activity.^20,24,56,57^ Nevertheless, in some cases the overlap with the experimentally known conformational change achieved via single residue perturbations is low, indicating the need to extend the analysis to multiple force insertions which is accomplished by MPR. As an application of the method, we focus on the position of maximizer residues in proteins selected to construct the sensing domain in GEFBs, in the context of relaying the conformational change to the reporter (fluorescent) domain(s). In these systems, the ultimate effect of ligand binding is a change in the chromophore environment of the FPs, which is responsible for its spectral output. In previous applications of PRS, the coordinates of either *apo* or *holo* structures have been used in constructing the **H** matrix, depending on the desired application. All the sensors in our selection are ‘direct response’ sensors, *i*.*e*., the fluorescence response is increased when the ligand binds. Therefore, we chose to identify the residues that play a significant role to drive the *apo*-to-*holo* or OFF-to-ON transition. In other words, we identify multiple residues in the *apo* conformation which, when perturbed in the optimized directions, will lead to a high overlap with the experimental/predicted *holo* coordinates. It should be noted that the protein is coarse-grained as a network of *N* nodes where any two nodes interact within a cut-off distance of 12 Å. Therefore, residues identified with our analysis pinpoint to approximate regions in the protein and should not be interpreted as exact residue indices.

### MPR solutions display positional and overlap convergence

The results of application of multiple forces on the selected cases up to five insertion points (*k* = 1-5) are presented in **Table 2** where the maximum overlap (*O*_max_) for each case and the residue indices leading to this result are listed. In all systems, increasing the number of perturbed residues display a convergence to the locations of the force insertion residues which we term ‘maximizer residues’ as the *O*_max_ values also exceed 0.8. However, the weights of the force vectors (equation 9) that drive that transition are significantly higher for KbP and CaM-M13 pairs both of which undergo a large domain reorganization, as indicated by large scaling factors applied when visualizing the force directions **(Table 2)**. In all systems, *O*_max_ values tend to converge after an initial jump from single to double perturbation except for MbP which already has a high *O*_max_ at *k* = 1. This is because the hinge bending motions are driven by a dominant eigenvalue that is easily triggered by single residue perturbations on many residues, as we have shown in detail in our original PRS studies.^20,21^

**Table 2.** Results for multiple force application on *k* residues*.

| Protein | $O_{max}$ / Maximizer Residue Indices | | | | | Vector Scaling Factor** | Sensor type / Insertion sites*** |
| --- | --- | --- | --- | --- | --- | --- | --- |
| | $k = 1$ | $k = 2$ | $k = 3$ | $k = 4$ | $k = 5$ | | |
| **KbP** | <u>0.74/</u><br>140 | **<u>0.88/</u>**<br>**17,97** | <u>0.89/</u><br>17,23,147 | <u>0.90/</u><br>17,30,61,139 | <u>0.91/</u><br>17,23,61,118,140 | 20 | Single FP/<br>1,151<sup>38-40</sup> |
| **CaM-M13** | <u>0.70/</u><br>109 | <u>0.80/</u><br>25,173 | <u>0.83/</u><br>37,125,173 | **<u>0.84/</u>**<br>**8,23,60,176** | <u>0.85/</u><br>6,23,60,129,173 | 20 | Single FP/<br>1,181<sup>42</sup> |
| **mICNBD** | <u>0.76/</u><br>120 | **<u>0.83/</u>**<br>**11,124** | <u>0.86/</u><br>11,55,124 | <u>0.88/</u><br>11,55,121,124 | <u>0.89/</u><br>11,55,117,124,125 | 10 | FRET/<br>1,126<sup>45</sup> |
| **MbP** | <u>0.94/</u><br>37 | <u>0.95/</u><br>21,330 | **<u>0.95/</u>**<br>**20,174,330** | <u>0.96/</u><br>20,174,300,330 | <u>0.97/</u><br>23,58,174,300,330 | 5 | Single FP/<br>175, 311<sup>11</sup> |
| **CitAP** | <u>0.63/</u><br>133 | <u>0.77/</u><br>101,132 | **<u>0.82/</u>**<br>**3,101,133** | <u>0.85/</u><br>3,101,131,132 | <u>0.88/</u><br>5,101,103,131,133 | 3 | Single FP/<br>1,133<sup>48</sup> |

\* Residue groups listed in bold are visualized in Supplemental Morph Movies S1-S5
\*\* Linear response dictates we can rescale the vectors proportionally to get the same overlap values; this factor is used in visualizing the vectors in reasonable size in Figures 1-4 and S2.
\*\*\* ‘Sensor type’ refers to either single fluorescent protein based, or FRET based sensors

In **Figure 1**, we display the detailed findings for KbP, along with a comparison of the performance of our two MPR approaches. In this protein, the predicted RMSD between the potassium loaded and free forms exceeds 17 Å **(Table 1)**, due to a major conformational change where the C-terminal BON and the N-terminal LysM domains pack against each other upon K^+^ binding, coordinated by the residues shown in **Figure 1A**.^49^ A major part of this transition may be achieved by perturbing a single residue on the mobile BON domain (P140, *O*_*max*_ = 0.74) **(Figure 1C and Movie S1)**. At *k* = 2, there is a significant jump in the *O*_*max*_ to 0.88 and the two maximizer residues are located on opposite domains with A17 on LysM and Q97 on BON. As *k* is further increased, there is an incremental change in *O*_*max*_ whereby we find that A17 is fixed as a maximizer residue while various residues near the C-terminus of the LysM domain appear as additional controllers of the conformational change (Y139, P140 or P147). A17 occupies an unconventional position for allosteric control, since it is in the middle of a long and flexible loop. However, it is on a loop supporting the potassium coordinating residues F6, V7, D9, A10, I77 and I80, and is 10.7 Å from the K^+^ ion, making it the perfect allosteric controller (**Figure 1A**). We note that for *k* > 2, D23 or K30 residing on the α-helix following A17 appears with nearly parallel force required to achieve the conformational change with high overlap. G61 at the region adjoining the two domains appears for *k* ≥ 4. In general, the main conformational change is controlled by A17 and one residue near the C-terminus; the forces on all other residues act as small corrections to these main drivers of the change.

**Figure 1.**
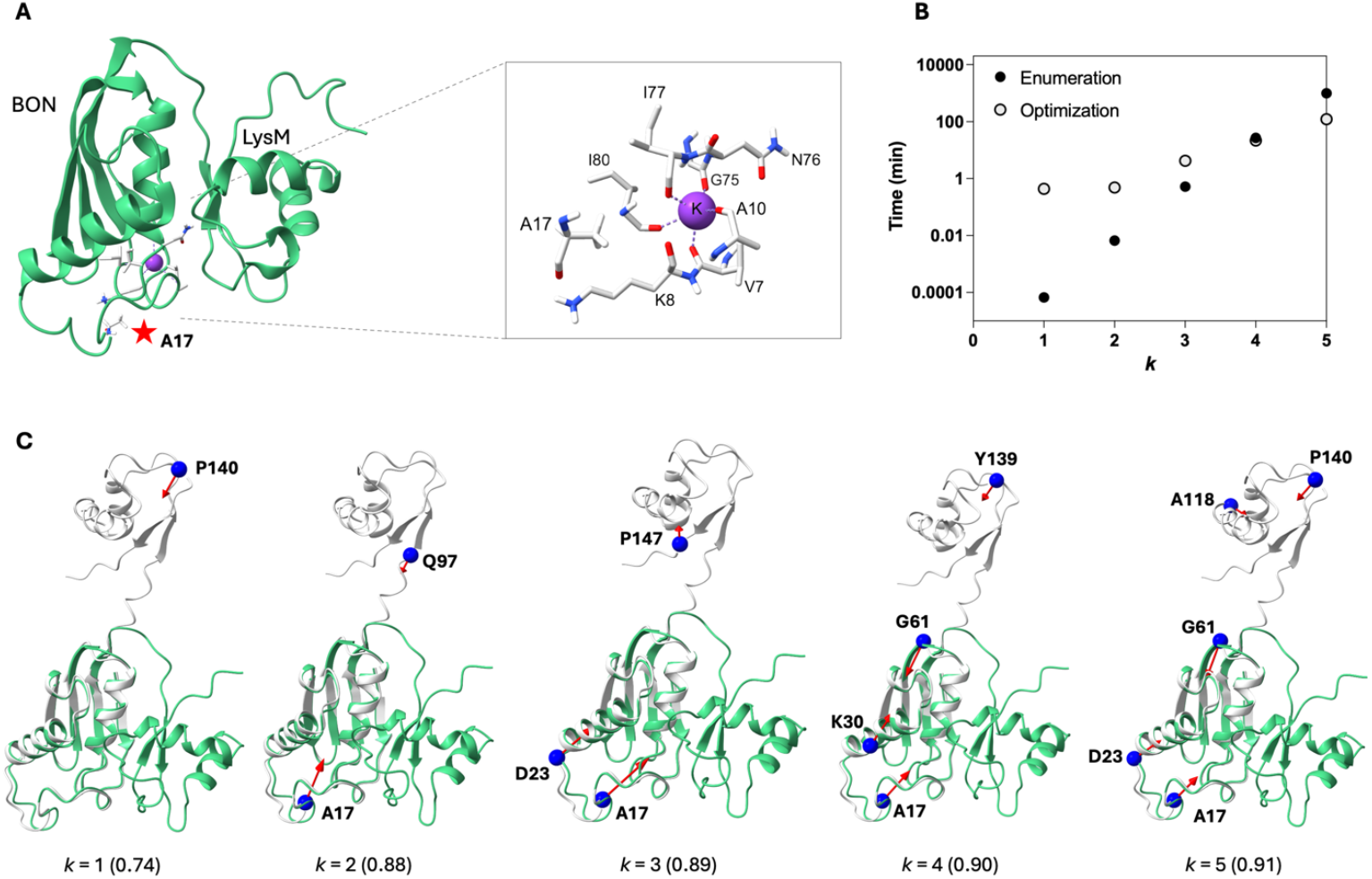
**A**. Structure of *holo* KbP highlighting the domains and potassium coordinating residues. **B**. The time required for the multiple force application with the two different approaches developed in this work, plotted on semi-log scale. **C**. Overlap maximizer residues in *apo*-to-*holo* transition of KbP for *k* = 1-5. Starting from *k* = 2, two residues on the opposite domains are located by MPR, and additional residues that appear at higher *k* are located near these residues (*apo* form in gray, *holo* form in green).

Convergence of perturbation locations for increasing *k* is observed for all the proteins studied, irrespective of the *O*_max_ values **(Figure S2)**. In general, *k* = 1 locates a residue on one of the domains. At *k* = 2, two residues on opposite domains are found to accomplish the conformational change with high overlap. For larger *k*, at least one of these two positions from the *k* = 2 solution is maintained while additional residues appear at locations close to these on either domain, reinforcing the effect. On the other hand, the vicinity of the original single residue from the *k* = 1 solution may (as in KbP, mlCNBD, CitAP) or may not (as in CaM-M13 and MbP) persist. In general, we suggest that rather than *O*_max_ convergence, residue location convergence be considered in deciding on the mechanism of allosteric control and to therefore scan the protein with MPR method for up to at least *k* = 4.

We demonstrate the computational cost of the calculations using the two approaches on KbP in **Figure 1B**. The enumeration approach displays a log-linear dependence on *k*, while optimization displays a slower increase in computational time. Hence, up to the break-even point of *k* = 4, enumeration is faster at solving for the maximizer residues, although both approaches complete the task within minutes. However, optimization significantly outperforms enumeration at *k* = 5 and is the only approach that leads to the findings in a reasonable compute time. For KbP with 151 residues, the analysis for *k* = 5 is completed in approximately 2 and 16 hours with the optimization and enumeration approaches respectively, on a single node of 112 CPUs with Intel(R) Xeon(R) Platinum 8480+ 2.0GHz and 224 GB memory.

### MPR justifies chain termini as insertion locations in single FP nanobiosensor designs

In single FP nanobiosensor designs, the operating principle is usually to insert the sensing domain at the bulge region of a circularly permuted FP (cpFP) adjoined by a suitable linker (**Figure 2A**). The constructs are optimized by a trial-and-error procedure to yield the maximum fluorescence response upon analyte binding.^11,44,58^ In theory, allosteric locations that convey the conformational change that occurs upon ligand binding to the bulge region are ideal insertion points. In practice, most designs utilize the two termini of the sensing domain protein as the starting point and make a high throughput generation of thousands of designs by combining cpFPs and the linkers from suitable libraries.^47,59^ Three of the five systems studied in this work are indeed such single FP nanobiosensors (KbP, CaM-M13, and CitAP; **Table 2**). We have presented the results of KbP in detail in the previous subsection which locates the C-terminus as an allosteric controller while the loop containing A17 also emerged as an allosteric controller, although it has not been utilized in GEFB designs. Below we refer to the other two systems.

**Figure 2.**
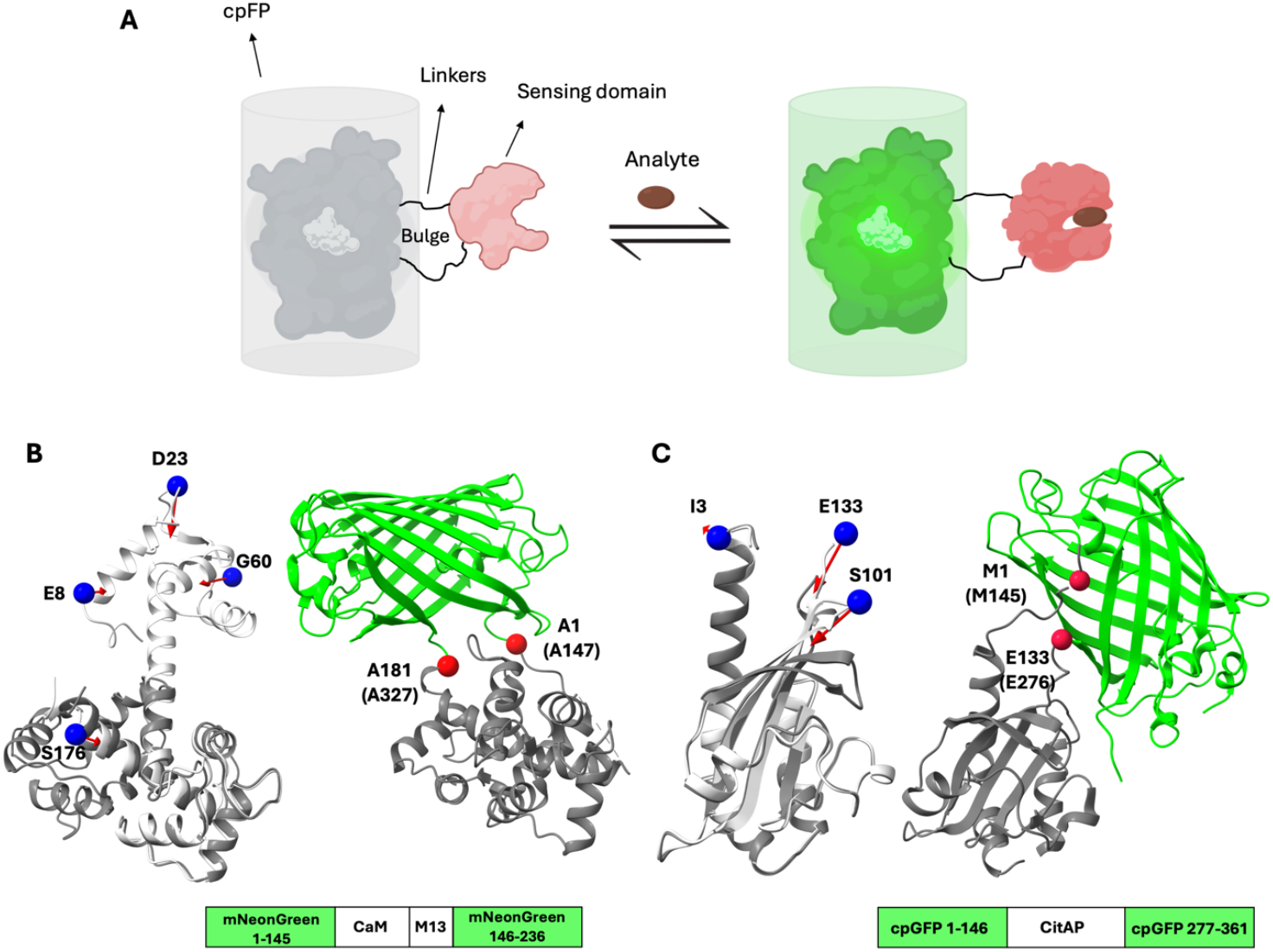
**A**. The general construct of single FP designs fuses a cpFP whose N- and C-termini are engineered at the bulge location near the chromophore with the protein selected as a sensor. Upon analyte binding, the fluorescence increases in direct response GEFBs depicted here, while the reverse is true for inverse response systems (created with biorender.com). **B**. Converging maximizer residues in CaM-M13 and their use as insertion points in fluorescent biosensors for NCaMP7 (PDB: 6XW2). **C**. Converging maximizer residues in CitAP and the insertion points in the CitrON sensor (PDB: 6LNP). Maximizer residues are shown as blue spheres on the *apo* (silver-initial) structure and superimposed with *holo* (dark gray-final) structure. Sensing domains in *holo* states are shown in dark gray and fluorescent domains are colored according to their emission. Insertion points are shown as red spheres. Residue labels in parentheses correspond to their indices in the reported sensor sequence.

CaM-peptide moiety is the most common sensing domain utilized in GEFB designs;^37^ however, in most cases, the peptide is covalently attached to the cpFP and non-covalently interacts with CaM. NCaMP7 is one of the few examples where the peptide is fused to both CaM and FP domains (see the construct in **Figure 2B**). CaM-M13 fusion exhibits a large conformational change where the central long helix is bent and the N terminal domain wraps onto the M13 peptide (**Movie S2**). We note that the initial structure in this case is a model that shows similarity to a low-calcium sensor structure and may not reflect the ‘true’ *apo* state (See **Preparation of PDB Files**). Perturbing a single residue, V109, on the fixed C terminal domain initially achieves an *O*_*max*_ of 0.7 and this value jumps to 0.8 when D25 and G173 are perturbed as a pair (**Figure S1D** and **Table 2**). Convergence in residue insertion locations is achieved at *k* = 4 whereby *O*_max_ = 0.84. Of the four residues found, D23 and G60 are on two EF-hand motifs in the N-terminal domain of CaM which bind Ca^2+^. We note that in our previous work utilizing PRS, we have determined the first EF-hand motif to be an important manipulator of CaM conformational change.^60^ In our later work, we have shown that turning off one charge on this motif suffices to significantly shift the conformational landscape of CaM^61^ cementing the allosteric role of this motif in CaM dynamics which was explored biochemically earlier.^62^ However, for the purposes of sensor design, these positions would be excluded as FP insertion points since they are also expected to hamper the binding of the targeted analyte Ca^2+^. On the other hand, the vicinity of G173 on the M13 helix seems to be a key location in manipulating the conformational change since it is conserved for all *k* > 1. In fact, the M13 helix or other peptides of similar length, is almost always the point of attachment of FPs in various optimized calcium sensor designs.^9,51,63^ For *k* = 4/5, E8/T6 emerge as the next position to trigger the conformational change near the N terminal tail where the FP mNeonGreen is fused in the NCaMP7 sensor. Thus, the positions implicated by MPR are in accordance with actual sensor designs which utilize a huge trial-and-error effort to optimize fluorescence response by conveying the signal generated by the conformational change while keeping the analyte binding region intact.

The citrate sensor CitrON is another example where FP insertion utilizes the termini of the sensor domain protein CitAP (**Figure 2C**). In CitAP, the main motion is the bending of the two β strands and folding of the connecting loop onto the citrate (**Movie S3**).^64^ Our analysis initially points to the C terminal residue E133 at *k* = 1, albeit with a relatively low *O*_*max*_ of 0.63. However, increasing to two perturbation points (*k* = 2) leads to a jump in *O*_*max*_ to 0.77, combining S101 on the ligand binding loop along with another C-terminal residue for conformation control. At *k* = 3, the N terminal residue I3 is included with another jump in *O*_*max*_ to 0.82. Adding more forces introduces incremental changes to *O*_*max*_ and the solutions keep suggesting residues on both C and N termini and the citrate binding loop (**Figure S2D, Table 2**). In CitAP, an overlap greater than 0.8 is achieved only when both N and C termini as well as S101 on the ligand binding loop are perturbed. As stated above for CaM-M13, it is best to avoid ligand binding residues as insertion points for the reporter domain. Therefore, the MPR suggested positions near the two termini are in accord with the experimentally optimized sensor design shown in **Figure 2C**, and in fact, with ligand binding acting directly on S101, constitutes the perfect allosteric communication scenario.

### MPR suggests insertion positions for FRET-based sensors

FRET is a physical phenomenon that occurs when two fluorescent molecules (a donor and an acceptor) are in proximity (up to 10 nm) and when the emission spectrum of the donor overlaps with the excitation spectrum of the acceptor. This principle is widely utilized in designing biosensors (**Figure 3A**). As in single FP designs, the donor and acceptor are attached to two domains of a sensing protein that move relative to each other in response to analyte binding. One such system is included in the set we study in this work where the cyclic nucleotide-binding domain (mlCNBD) of bacterial M*Loti*K1 channel is employed (mlCNBD-FRET; **Figure 3B**). The main motion during the *apo*-to-*holo* transition involves a large displacement of the N- and C-terminal helices, while the ligand-binding site remains relatively immobile (**Movie S4**). The movement of the C-terminal helix is part of a larger regulatory mechanism; the nucleotide-bound CNBD interacts with the transmembrane domain of the MLotiK1 channel in a fourfold symmetric arrangement, causing the channel to gate.^65^ Such a large conformational change at distant sites makes this protein an ideal sensing domain for nanobiosensors and also for our investigation of allosteric control.

**Figure 3.**
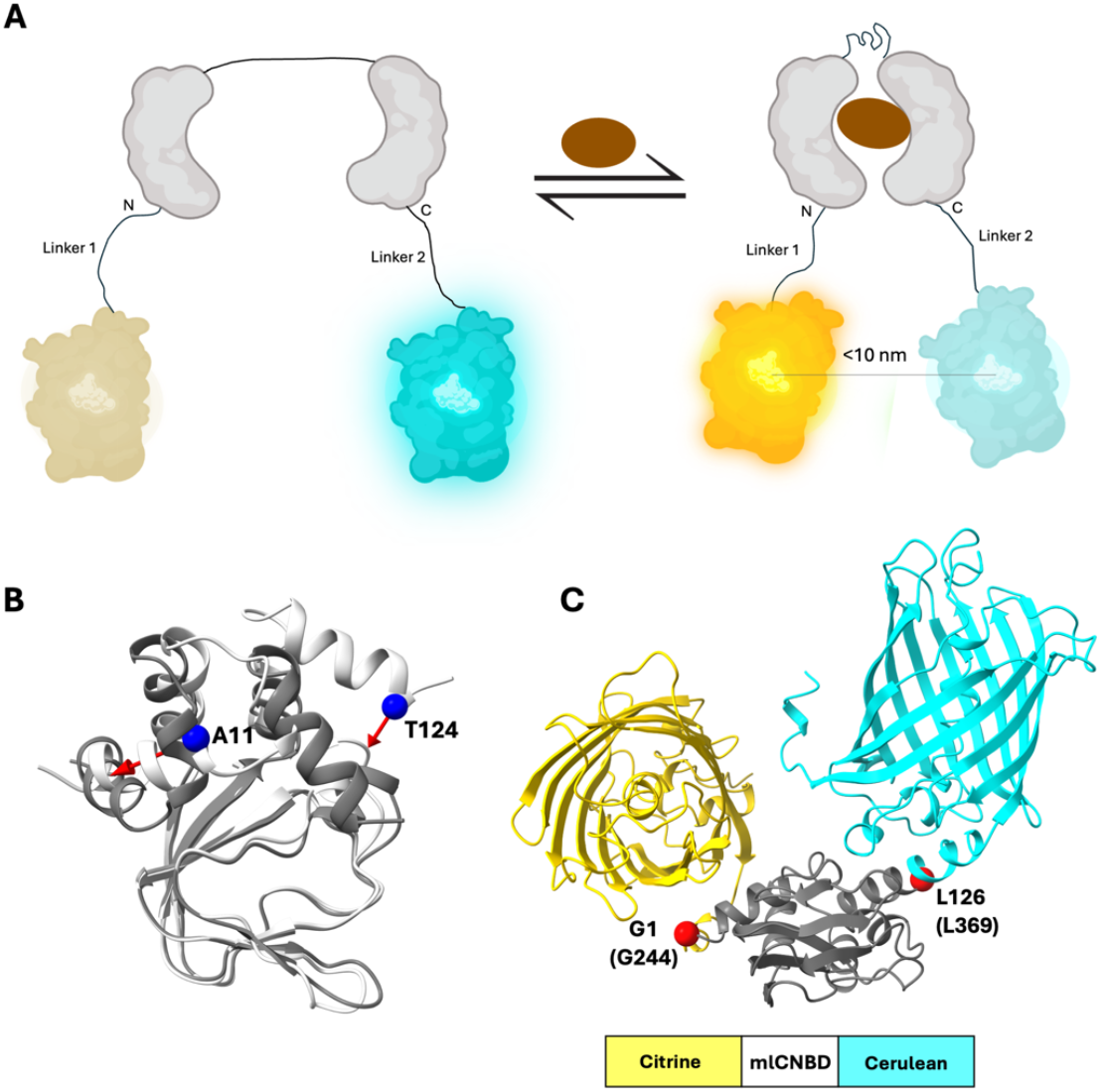
**A**. The general construct of FRET-based designs fuses two FPs with the protein selected as a sensor; upon analyte binding, the FPs are expected to get closer and give a response due to FRET (created with biorender.com). **B**. Converging maximizer residues in mlCNBD and **C**. their use as insertion points in the corresponding FRET sensor (structure is predicted with AF3 server, see Methods). Color and numbering scheme are the same as in Figure 2.

In mlCNBD, our analysis initially points to residue I20 near the C terminus at *k* = 1 with an *O*_*max*_ of 0.76. At *k* = 2 a jump in *O*_*max*_ = 0.83 occurs whereby perturbing the A11/T124 combination captures a significant portion of *apo*-to-*holo* transition. Increasing *k* further leads to incremental increase in *O*_*max*_ and the maximizer residues always include this pair (**Table 2**). In general, broadly three regions are suggested: residues on the C terminal helix (115-126), A11 on the N terminal helix and V55 which sits on a β strand that is a non-mobile hinge point around which the helical bundle rotates (**Figure S2B**). The two ends of this protein were used to attach two FPs to create the FRET sensor in conformity with the regions pointed by MPR **(Figure 3B**).^45^ We note that inserting a single circularly permuted FP (cpFP) at a loop near the cAMP ligand (P65-N66, P285/N286 according to numbering in PDB 1VP6) also resulted in an efficient sensor (G-Flamp1^44^), however our analysis did not indicate that region as an allosteric hotspot. In fact, in G-Flamp1, the mere presence of the cAMP in the ON state effects the conformation of nearby residues which interact with the chromophore. Therefore, its main activation mechanism depends on the proximity of the ligand to the chromophore rather than an allosteric conformational change at a distant site.

### Prediction of allosteric insertion sites on non-terminal regions by MPR

While most single FP nanobiosensors rely on terminal site selection followed by optimization of cpFP and linkers, there have been attempts at determining allosteric locations for their guided rational design mostly using periplasmic binding proteins as sensing domains.^11,15,66,67^ One such case is presented by the maltose biosensor whereby a hinge bending motion dominates the conformational change of the sensor domain protein MbP (**Movie S5**). We note that FRET sensors utilizing MbP have also been constructed (FLIPmal^46^). However, these fuse the two FPs to the N- and C-terminus of MbP and seem to make use of the hinge motions triggering the closing-in of the two lobes. Considering the relatively large size of MbP (370 residues) and the distance requirement of donor/acceptor in FRET sensors, we conclude that these do not utilize allosteric signals but rather make direct use of the large intrinsic motions for efficiency.

In designing the single FP maltose sensor, the authors have hypothesized that coupling ligand binding to fluorescence requires that the insertion site of cpGFP must transmit global conformational changes of MbP to the cpFP chromophore.^11^ Structural analysis therein showed significant torque at non-terminal residues N173 and L311 upon maltose binding (designated as MBP175-cpGFP.L1-HL and MBP311-cpGFP.L2-NP in the original paper), suggesting these changes could trigger maltose-dependent fluorescence.^11^ This is therefore one of those few constructs where the fusion is achieved at non-terminal chain location in the protein. In this case, the FP is sandwiched between the MbP in the construct (**Figure 4**), whereby insertion at either of these two locations provided detectable fluorescent signals. Their structures have been solved by X-ray crystallography (PDB codes 3OSQ and 3OSR, respectively for the position 173 and 311 constructs).

**Figure 4.**
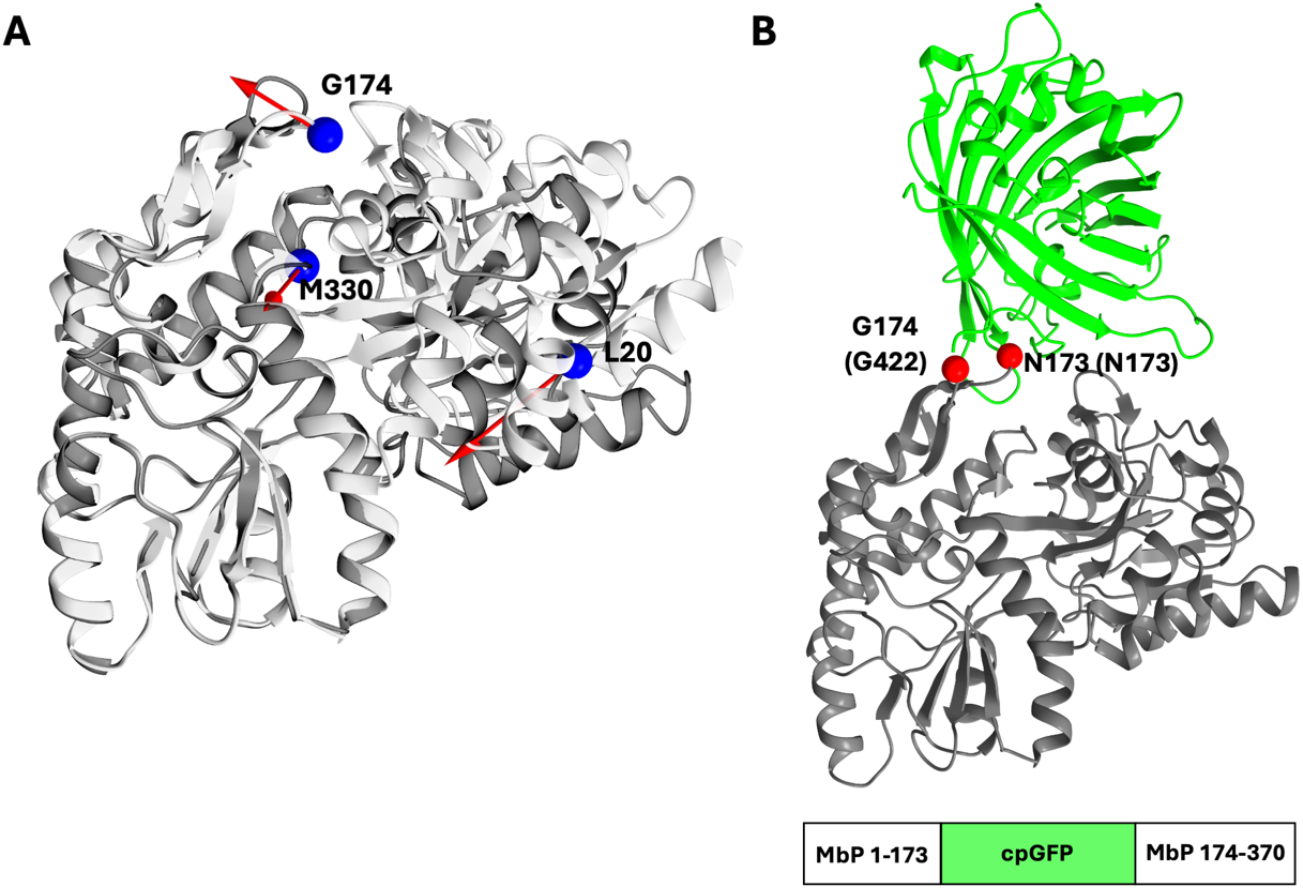
Converging maximizer residues in the **A**. MbP protein and **B**. their use as insertion points in the corresponding maltose sensor (PDB:3OSQ). Color and numbering scheme are the same as in Figure 2.

We therefore set out to search the perturbation sensitive sites disclosed by MPR in this protein. In MbP, the *apo*-to-*holo* transition which is a hinge bending motion from closed-to-open state, could be achieved by perturbing only one residue (V37, *O*_max_ > 0.9, **Table 2**). This result is consistent with the previous observation where for hinge-bending proteins such as MbP and ferric binding protein A, a high overlap could be achieved by single residue perturbations.^21^ At higher *k* values positional convergence is achieved. M330 which is in direct contact with the sugar appears at all *k* (**Figure S2C**) along with L20, A21 or V23 which are in direct contact with V37. G174 comes into the picture with *k* = 3, which is the position equivalent to the insertion location in the construct mentioned above (**Figure 4**). We also find the region constituting the hydrophobic patch between L20/V21/V23/V37 to be an allosteric modulator of MbP, although no construct utilizing this position has been synthesized or tested.

This sensor provides an excellent example for guiding rational design. In their work Marvin *et al*. first resorted to typical metrics used for quantifying local structural perturbation, such as solvent accessible surface area or distance metrics, but these did not yield satisfactory results.^11^ They then analyzed the changes in the dihedral angles spanning four consecutive C_α_ atoms to identify residues which were significantly torqued upon ligand binding to arrive at candidate locations for the design. Such approaches require close monitoring of the structural changes on a case-by-case basis to identify residues of interest. Here, we show that MPR provides an alternative approach, blind to the mode of the binding or the size of the conformational change to identify regions of interest.

## Conclusions

In this work, we utilized multiple perturbed responses (MPRs) in proteins to identify hotspots in small molecule binding proteins. Using an optimized approach, we were able to reveal multiple allosteric hotspots that are key in driving the *apo*-to-*holo* transitions with a reasonable computational power. These residues tend to cluster on two or three separate regions on the protein, with some of them being on mobile regions while others acting as fixed hinges. The approach relies on linear response theory and expands on our previous work that relates responses triggered by single residue perturbations with allosteric regions. MPR may find many uses within the framework of allosteric control.

Here we considered GEFBs as model systems to test the validity of our results and we present case studies illustrating the applicability of the method to predict hotspots in sensing domains. Regions identified by PRS were found to be critical for transmitting the conformational change triggered by ligand binding in previous work. As GEFBs operate on allosteric coupling between the ligand binding site and the FP, we considered forces inserted on residues that maximize the overlap with the conformational change due to the *apo*-to-*holo* transition. Our analyses revealed some dynamically important regions that were used in real biosensor designs and suggested some additional sites that have not been reported to date. It is worth noting that which locations to choose from depends on experimentally imposed constraints since insertion points may not all be possible to engineer. Some residues suggested by MPR may not be optimal since they are on key secondary structural elements which would compromise the structural integrity of the protein. They may also be at an immobile region which would not change the environment of the chromophore significantly when the ligand binds, leading to a non-responsive sensor. In some cases, the maximizer residues are directly at the ligand binding sites (as in CaM-M13 and CitAP) and therefore would not be preferred as insertion sites. Depending on the application, our code can be modified to exclude such positions to be excluded from the solution, although we refrained from doing that in this work to analyze the total output.

In four of the five systems we studied, both the FRET and single-FP sensor constructs utilized the two termini of the sensor domain protein as insertion sites for the FPs. While this selection makes sense in terms of optimal experimental design, MPR indeed suggested residues near one or both termini as allosteric spots in all these cases, justifying the allosteric control imposed by these regions in the success of conveying the conformational change to the reporter domains. However, in those cases where the NPR suggested sites are near the termini (e.g. residue 11 in mlCNBD, or residue 3 in CitAP), one might consider truncating some of the extra residues to get improved fluorescence response.

Our approach of using allosterically relevant sites can be useful for cases when insertion of a reporter domain at a particular site introduces steric impediments to achieving the ligand binding-induced conformational change. For example, the flexible hinge regions, which are commonly used as insertion points in periplasmic binding proteins are shorter in some members and may not tolerate the insertion of FP domains as they might hamper folding or ligand affinity.^68^

In conclusion, MPR offers a computationally effective approach to reveal multiple (allosteric) sites that may be functionally relevant in biosensor design. The examples we present show that the number of force insertion points may differ depending on the target system. To get an insight on which regions may be viable as FP insertion sites, we recommend scanning the protein up to *k* = 5, which is achievable within a few hours for a protein of ∼150 residues using the optimization approach and focusing on the convergence of the locations pinned by MPR.

One limitation in applying the PRS approach has been the requirement of having experimental structures for both conformational states of the protein. In this work, we have also proposed a protocol to predict *apo* and/or *holo* structures of the proteins of interest by tweaking the parameters of AF2 algorithm, in case of missing experimental structures. There is a growing number of studies which recommend using a shallower MSA with a smaller number of aligned sequences as a means to increase the conformational diversity of the predicted models and our results support that trend.^69–72^

We would also like to emphasize that the positions suggested by MPR depend on the Hessian constructed using a preselected cutoff distance, 12 Å in this work. We utilize **H**^-1^ as an approximation to the **C** matrix. For more precise predictions, we suggest carrying out MD simulations under conditions where the sensor will be utilized, e.g. the ionic strength and pH of the working environment. 20-40 ns chunks of equilibrated MD simulations provide excellent predictors of the **C** matrix (equation 1) via the calculation of the variance-covariance matrix and is free of the problem of cutoff distance while also including environmental effects.^73^

Another research area where MPR may be used is in determining physics-based reaction coordinates (RCs) for exploring the conformational landscape of a protein. The area of collective variable selection is actively explored^74^ with many works relying on the principal components (PCs) or time dependent components (tICs)^75^ as the RCs on which to project the conformations. The advantage of selecting physics-based RCs is that, when comparing the same system under slightly modified conditions, the differences may always be followed intuitively and comparatively. Conversely, when using PCs or tICs, there is no guarantee that the projections reflect similar phenomena, and this renders interpreting shifts in the energy landscapes tricky. In fact, even when analyzing two MD simulations for the same system under the same conditions, one might get completely different energy surface projections.^76^ To prevent this situation, we have previously shown that a PRS determined single residue coupled with a distant fixed point would facilitate the conformational transitions in steered MD simulations^77^ and this approach would later discriminate the conformational preferences of ligand bound vs. ligand free phosphorylated RAS.^78^ Building upon the success of the single residue perturbation locations, we propose that with the development of MPR, the *k* = 2 solution may now be used as a natural CV in advanced sampling techniques. We believe that MPR approach we developed in this work presents itself as a general, computationally efficient tool for exploring allosteric phenomena in protein dynamics.

## Code availability

Codes calculating the displacement vector, solving the *O*_max_ and creating .bild files to visualize force vectors in ChimeraX are provided in https://github.com/midstlab/MPR. A sample solution is also provided for KbP up to *k*=5.

## Supporting Information

Movie S1. *Apo-*to*-holo* morph of KbP Movie S2. *Apo-*to*-holo* morph of CaM-M14 Movie S3. *Apo-*to*-holo* morph of CitAP Movie S4. *Apo-*to*-holo* morph of mlCNBD Movie S5. *Apo-*to*-holo* morph of MbP

## Acknowledgments

This work was financially supported by TUBITAK project no.121Z329. Python codes and Gurobi was run on TUBITAK ULAKBIM, High Performance and Grid Computing Center (TRUBA resources).

## Declaration of generative AI and AI-assisted technologies in the writing process

During the preparation of this work the authors used ChatGPT in order to improve the readability and language of the manuscript. After using this tool, the authors reviewed and edited the content as needed and take full responsibility for the content of the published article.

## CRediT authorship contribution statement. Melike Berksoz

Writing – review & editing, Writing – original draft, Visualization, Validation, Software, Methodology, Investigation, Formal analysis, Data curation. **Ali Rana Atilgan:** Writing – review & editing, Writing – original draft, Validation, Project administration, Methodology, Investigation, Formal analysis, Data curation. **Burak Kocuk:** Writing – review & editing, Software, Methodology, Investigation, Formal analysis. **Canan Atilgan:** Writing – review & editing, Writing – original draft, Supervision, Resources, Project administration, Methodology, Investigation, Formal analysis, Data curation, Funding acquisition, Conceptualization.

