## Supplementary material for "Multiply Perturbed Response to Disclose Allosteric Control of Conformational Change: Application to Fluorescent Biosensor Design"

**Figure S1.** Alphafold Models for **A.** KbP **B.** CaM-M13 **C.** CitAP. Top five models are displayed in the order of model accuracy ranking.

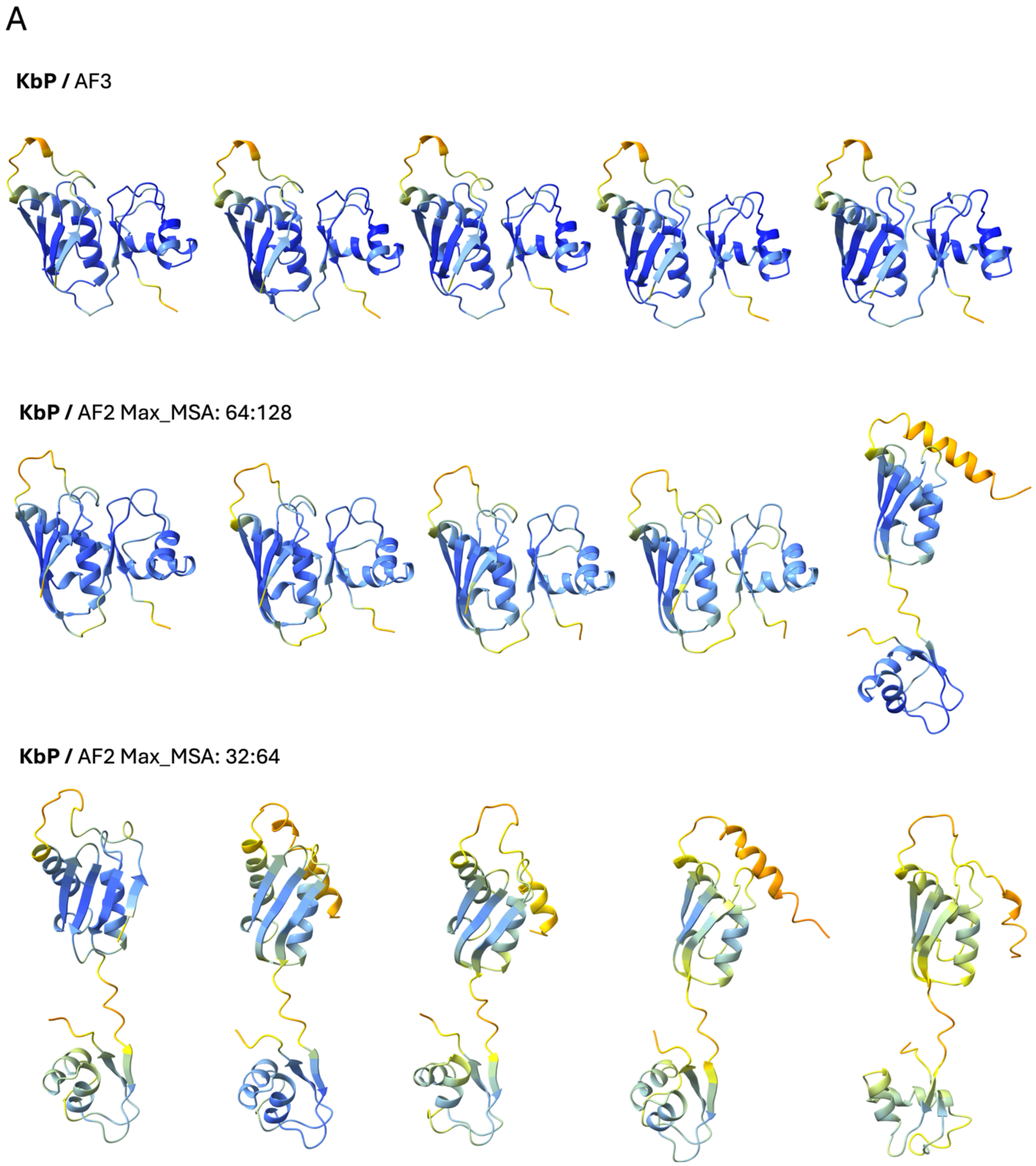

B

**CaM-M13 /** AF3

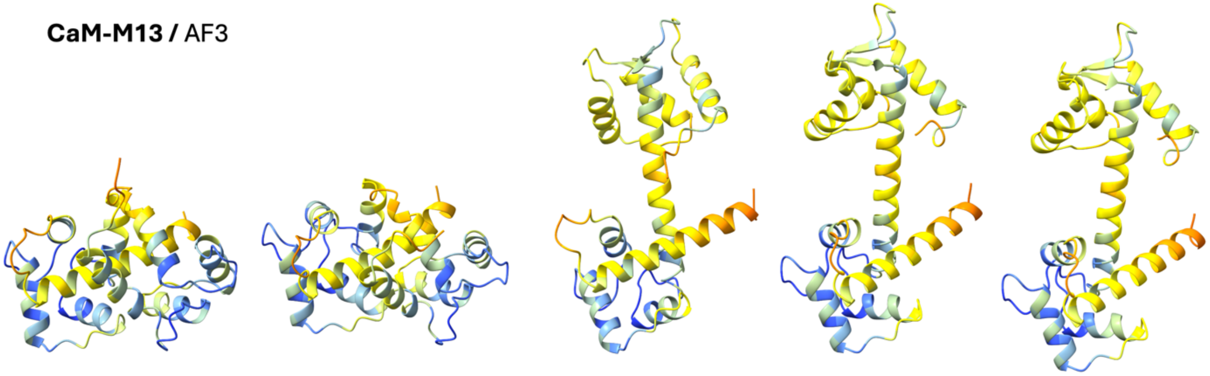

C

**CitAP /** AF2 Max_MSA: 32:64

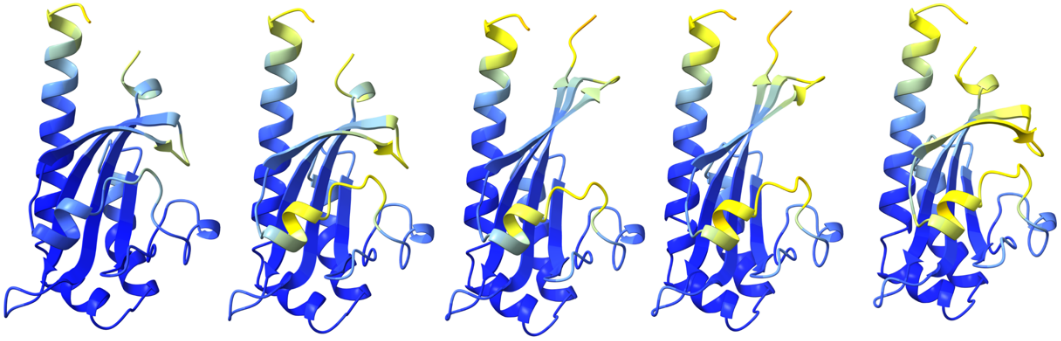

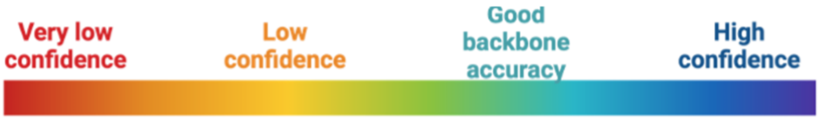

**Figure S2.** Maximizer residues for *apo*-to-*holo* transition of **A.** CaM-M13 **B.** mlCNBD **C.** MbP **D.** CitAP**.** Apo states are colored silver in each.

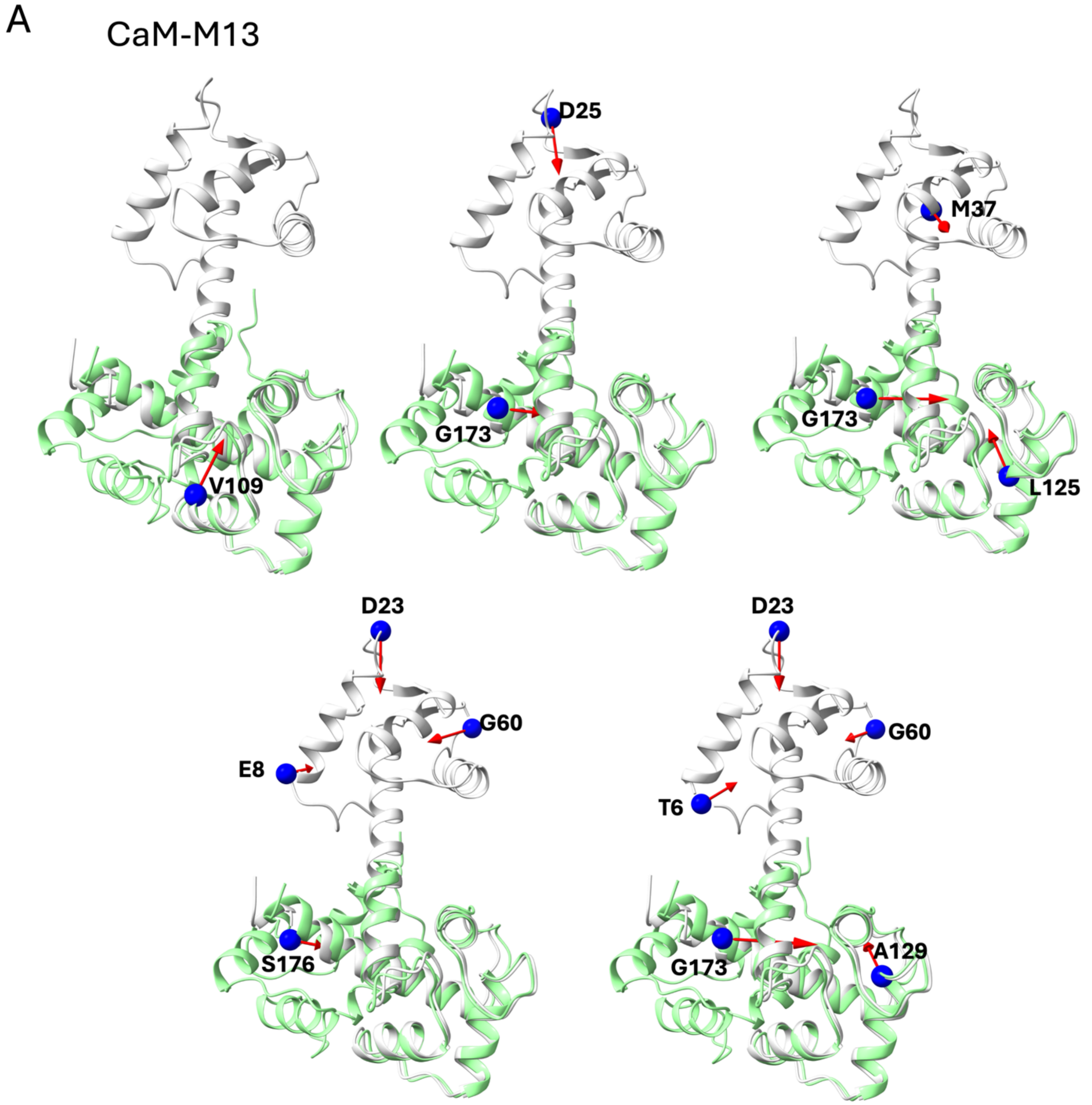

B

mlCNBD

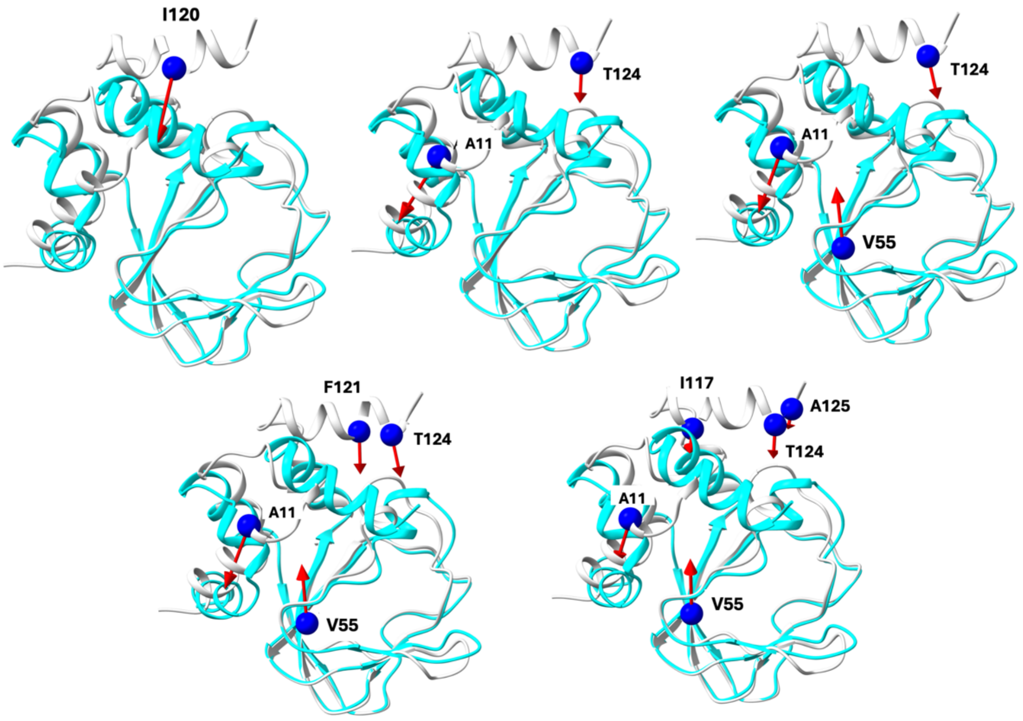


C

MbP

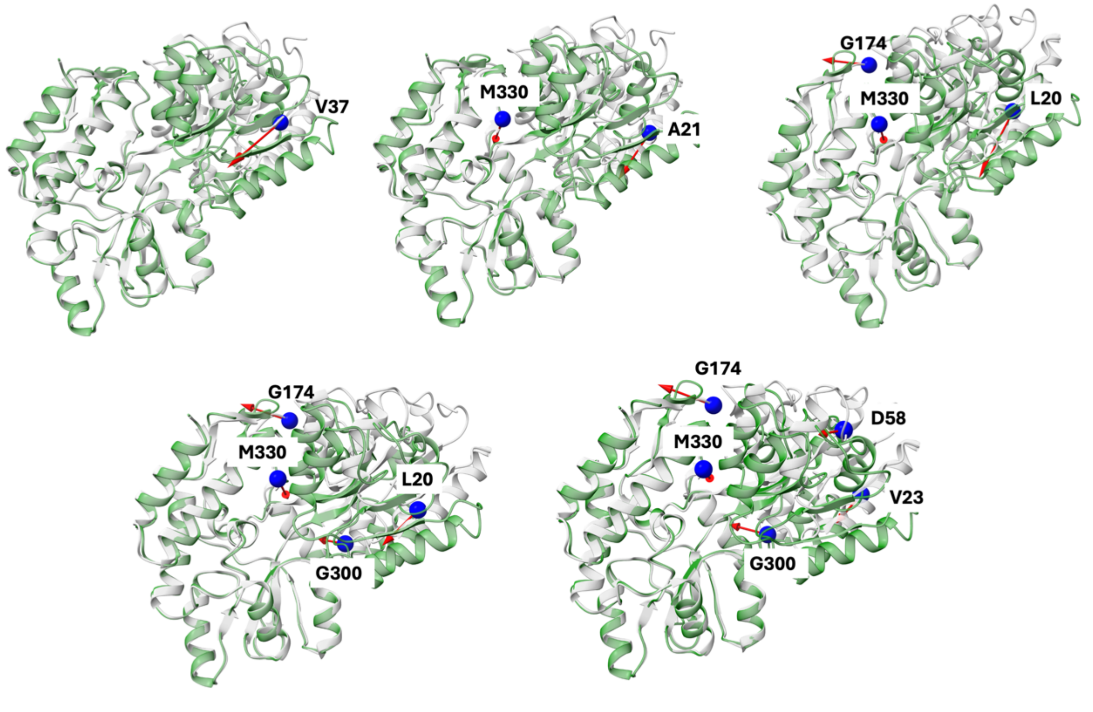

D

CitAP

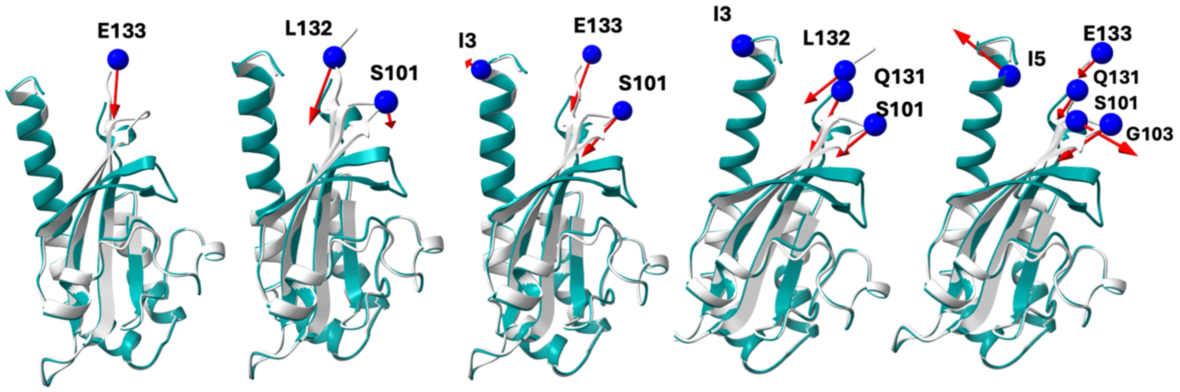
E133
L132
S101
I3
E133
S101
I3
L132
Q131
S101
I5
E133
Q131
S101
G103
